# Harvest probabilities of European ducks: the very first estimation based on reward rings

**DOI:** 10.64898/2026.02.20.706967

**Authors:** Souchay Guillaume, Caizergues Alain, Bacon Léo, Champagnon Jocelyn, Devineau Olivier, Gelin Marion, Grzegorczyk Emilienne, Lebreton Jean-Dominique, Plaquin Betty, Pradel Roger, Guillemain Matthieu

**Affiliations:** Service Santé de la faune et fonctionnement des écosystèmes agricoles, DRAS, Office Français de la Biodiversité - Nantes, France; Service Conservation et Gestion Durable des Espèces Exploitées, DRAS, Office Français de la Biodiversité - Nantes, France; Service Anthropisation et Fonctionnement des Ecosystèmes Terrestres, DRAS, Office Français de la Biodiversité - Pérols, France; Tour du Valat, Research Institute for the Conservation of Mediterranean Wetlands - Arles, France; University of Inland Norway, Department of Forestry and Wildlife Management, Campus Evenstad - Koppang, Norway; Service Conservation et Gestion Durable des Espèces Exploitées, DRAS, Office Français de la Biodiversité - Arles, France; Service Conservation et Gestion Durable des Espèces Exploitées, DRAS, Office Français de la Biodiversité - Villiers-en-Bois, France; CEFE, Montpellier University, CNRS, EPHE, IRD - Montpellier, France; Service Conservation et Gestion Durable des Espèces Exploitées, DRAS, Office Français de la Biodiversité - Ile d’Olonne, France

**Keywords:** Harvest, management, recovery, reporting, ringing, survival

## Abstract

Ringing-recoveries are an overarching element of population dynamic studies that allow estimating mortality causes and hence improve wildlife management. However, possible drawbacks of recovered rings reside in the fact that reporting probability is rarely known, but consistently lower than 100%. Thus, estimating harvest probabilities (mortality probability due to harvesting) of exploited species without knowledge of ring reporting probability by people exploiting these animals is not straightforward. We here provide the first ever reward-ring study carried out to evaluate European reporting probabilities, hence European harvest probabilities, in three species of ducks (Mallard *Anas platyrhynchos*, Eurasian Teal *A. crecca* and Common Pochard *Aythya ferina*). The 70 Euros reward on some rings was considered to yield a total return of the rings, allowing by comparison to evaluate the reporting probability of standard rings. After the initial year of ringing, annual reporting probability was very similar among the three species, at 0.70-0.76, suggesting two-thirds of the found rings are sent back to the ringing centre. This allowed computation of the annual harvest probability, which was up to 0.25 during the first months after ringing in fall but decreased to 0.04-0.09 during later years. Compared to North American results, the present estimates suggest birds are subjected to a heavy hunting mortality during the first months after ringing, but this pressure declines in later years, likely owing to counter selection of vulnerable/exposed individuals and/or learning by the birds.

## Introduction

Management of exploited populations increasingly relies on predictive demographic models, used to simulate the potential consequences of concurrent policy scenarios (e.g. Williams et al., 2002; Madsen et al., 2015). Among the best known examples, this is how adaptive harvest management of North American waterfowl is set-up (Nichols et al., 2007). The likelihood that individuals of a population will be taken over a period of time (termed *Harvest probability*) is a central parameter of such population dynamic models.This is an essential information to evaluate the level of exploitation in the population, the relative share of natural vs. human-induced mortality, and to assess the level of implementation of harvest policy in the field (e.g. Royle & Garrettson, 2005; U.S. Fish and Wildlife Service, 2006; Boomer et al., 2013; Arnold et al., 2020; Riecke et al., 2022).

Harvest probability is generally derived from the information provided by rings or tags returned by hunters/fishermen, but not all such marks are returned when found (Henny & Burnham, 1976). It is therefore not possible to derive harvest mortality/probability from ring recovered by hunters directly (the proportion of deployed marks later recovered by hunters and sent back to scientists). In that matter, knowledge about mark reporting probability (the proportion of marks recovered by hunter and sent to the marking scientists) is key (e.g. Pollock et al., 1995; Royle & Garrettson, 2005).

Several methods exist to evaluate mark reporting probability in exploited fish and wildlife populations, including questionnaires sent to users or secretive planting of marks in the bag or basket of already shot/captured animals and monitoring of their subsequent potential report (e.g. Geis & Atwood, 1961; Martinson & McCann, 1966; Martinson, 1968; Murphy & Taylor, 1991; Nichols et al., 1995). However, the most robust, precise and widely used method to do so is to use rings (for birds) or tags (for fish) associated with a reward (Bellrose, 1955; Henny & Burnham, 1976; Pollock et al., 1994, 1995; Nichols et al., 1995; Boomer et al., 2013). Such reward has to be of sufficiently high value to expect a 100% reporting probability of recovered marks (e.g. Conroy & Williams, 1981; Nichols et al., 1991; Royle & Garrettson, 2005; Zimmerman et al., 2009); the term “High-reward tag method” is commonly used in fisheries (e.g. Hoenig et al., 1998; Pollock et al., 2001; Meyer et al., 2012). Using this approach, mark reporting probability can simply be evaluated by the ratio of the recovery probability of standard marks to the recovery probability of reward marks, under some hypotheses related to independence of reporting probabilities between standard and reward marks, and a stable behaviour and good information of the people finding the marks (e.g. Pollock et al., 2001; Williams et al., 2002).

Reward mark schemes have been set-up on fish in a variety of situations (Murphy & Taylor, 1991; Hoenig et al., 1998; Pollock et al., 2001, 2002; Taylor et al., 2006, 2021; Meyer et al., 2012). In birds, the method has only been used in a handful of species, i.e. Wild Turkey *Meleagris gallapavo* (Difenbach et al., 2012), Ring-necked Pheasant *Phasianus colchicus* (Diefenbach et al., 2000), Mourning Dove *Zenaida macroura* (Tomlinson, 1968; Reeves, 1979; Scott et al., 2004; Sanders & Otis, 2012), and some waterfowl: Mallard *Anas platyrhynchos* (e.g. Bellrose, 1955; Henny & Burnham, 1976; Nichols et al., 1991, 1995; Reinecke et al., 1992; Royle & Garrettson, 2005; Boomer et al., 2013), American Black Duck *A. rubripes* (Conroy & Blandin, 1984; Garrettson et al., 2014), Wood duck *Aix sponsa* (Hine et al., 2004; Balkcom et al., 2010; Garrettson et al., 2014), Canada *Branta canadensis*, Cackling *B. Huthinsii*, Ross’s *Chen rossii* and Snow goose *C. caerulescens* (Zimmerman et al., 2009, 2010; Souchay et al., 2014). Such studies have highlighted differences in mark reporting probabilities between geographic regions (e.g. Nichols et al., 1995; Zimmerman et al., 2009; Boomer et al., 2013), changes in mark reporting probability over time (e.g. Murphy & Taylor, 1991; Arnold et al., 2020) and between species (Meyer et al., 2012), in line with the behaviour of hunters, the ease with which marks can be returned or the frequency at which such marks are encountered (e.g. Pollock et al., 2001; Garrettson et al., 2014; Souchay et al., 2014; Arnold et al., 2020).

Despite its importance for hunting sustainability and conservation, the impact of harvest on survival and hence the population dynamics of exploited waterfowl has seldom been explored in Europe (Devineau, 2010), with recurrent calls for a proper estimation of harvest probabilities and a widening of adaptive harvest management schemes (Elmberg et al., 2006; Holopainen et al., 2018). The European Commission is now pressing further, considering that it is the responsibility of those willing to harvest bird populations to evaluate the likely consequences of harvest beforehand (Samarelli & van der Stegen, 2024). Unfortunately, no reward ring programme such as the ones described above in North America had ever been implemented in Europe, and even in North America the number of species considered is still limited (see above). Consequently, the rare demographic models for ducks available in Europe had to rely on North American harvest probability estimates, sometimes obtained over a different period and a different species, despite the geographic, temporal and species-specific effects listed above (e.g. Devineau et al., 2010 for Eurasian Teal *Anas crecca* - hereafter Teal - relying on North American Mallard data).

The aim of the present paper is to fill this gap by providing ring reporting probability and hence harvest probability estimates through the first ever bird reward ring scheme in Europe, simultaneously for Mallard, Teal and Common Pochard (*Aythya ferina*, hereafter Pochard). Mallard numbers wintering in France have been stable over the long term (1980-2025) but have declined over the sort-term (2014-2025), Teal numbers have increased both over the long and short-terms, and Common Pochard numbers have declined over both time scales (LPO 2025). National hunting bags for Mallard and Teal during the 2023-24 hunting season were estimated at 695,000 and 300,000, respectively, while Common Pochard bag estimate was too imprecise to be published (Lapébie et al. 2025). Teal and Common Pochard are on the priority list for the European Commission owing to their poor conservation status in the EU and apparent unsustainable current harvest (Cruz-Flores et al., in prep.).

## Material and methods

### Capture and ringing

Ducks were captured using a variety of swim-in traps, vertical nets and drop-door traps depending on species and environmental conditions in 11 areas scattered over France, discarding sites where only a handful of birds could be captured (Figure 1): capturing and ringing enough ducks over extended periods of time (tens of years sometimes) is a tedious work, so the motivation of ringers can only be sustained when sufficient numbers of birds can be captured regularly. This marginally depends on local logistics (ease of access, number of available staff), but is mostly dependent on the number of wintering ducks locally. The ringing sites were therefore generally placed in major winter quarters for the given species (only Grand-Lieu lake in western France for Common Pochard in those years), which is generally associated with good hunting opportunities and hence often a relatively high local hunting pressure. However, these birds are very mobile even during the winter (e.g., Caizergues et al., 2011; Gourlay-Larour et al., 2012; Tableau et al., 2022), so they could be hunted and recovered anywhere, in a completely different area with a very different local hunting pressure. All parameters estimated here are annual, hence covering the whole annual cycle and migratory range of these species.

**Figure 1.**
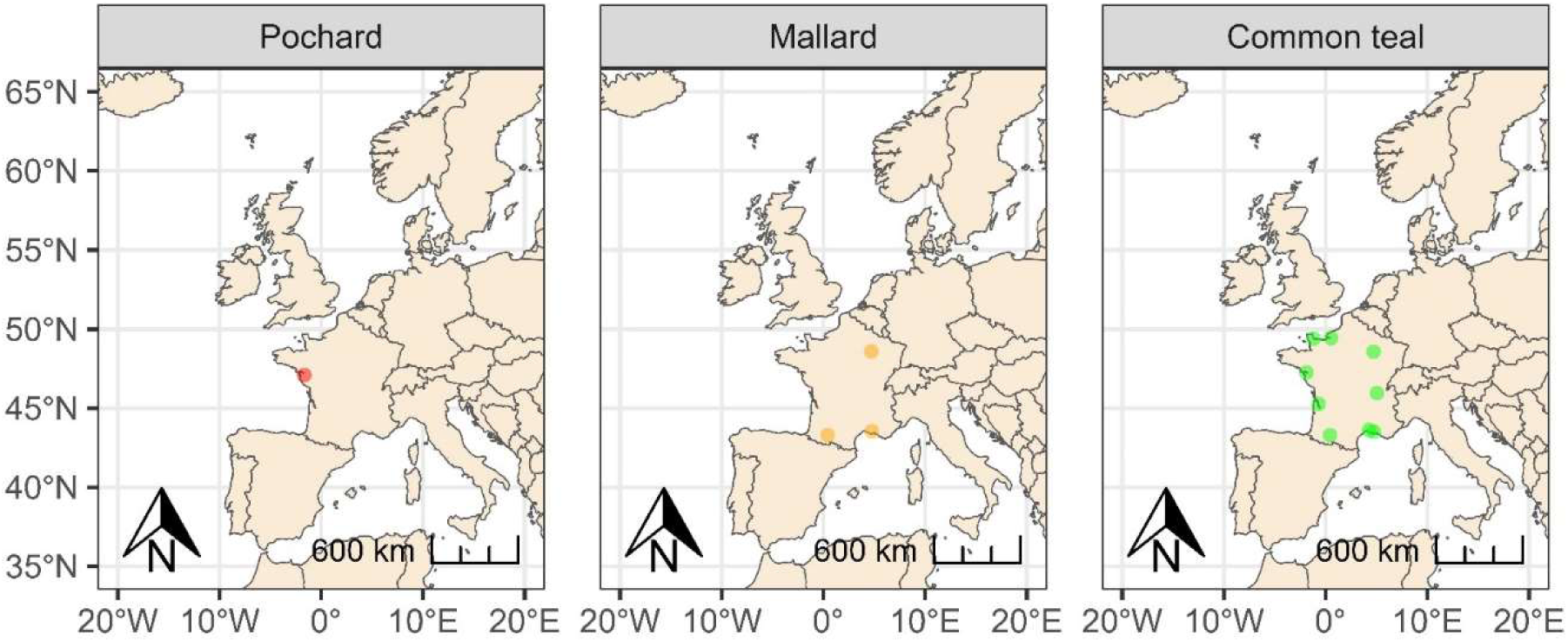
Map of the 11 duck ringing areas used to deploy standard and reward rings in the three species in France, 2011-2020.

Based on the number of catches during earlier years, it was initially computed that fitting 100 reward rings annually over three years on each species would allow estimating reporting probability adequately (Devineau et al., 2006). In practice catches were poorer than expected during the initial study years, so it took more years than expected and a total of 116/727 (Mallard), 252/3526 (Teal) and 184/992 (Pochard) reward and regular rings were respectively fitted onto birds from 2011 to 2020 (some recoveries occurring until 2021). In line with other research projects, a nasal saddle of the type described by Rodrigues et al. (2001) was fitted onto all Pochards and ca. two-thirds of the Teal (e.g. Guillemain et al., 2010; Tableau et al., 2022). Mallards were not saddled.

Absolutely no selection occurred in the field, and a reward ring was fitted randomly to every sixth to seventh captured duck of a given species, in addition to a regular numbered identification ring on the other leg, independently of sex and age (which were determined after plumage examination following Mouronval, 2016). Inscription on the regular ring only provides information on the national ringing center (here, the National Museum of Natural History in Paris), ring size coded as two letters, and a unique ID number, hence for example “MUS. PARIS FX153251”.

In order to avoid hunters possibly encountering many reward and standard rings locally, and therefore changing their reporting behaviour (Henny & Burnham, 1976; Pollock et al., 2001; Meyer et al., 2012), reward rings were scattered over the ringing sites whenever possible (except in Pochard where most ringing at national scale occurs at one single site anyway). Reward rings were identical to regular numbered rings, made of stainless steel and provided by the same manufacturer (I.Ö. Mekaniska AB, Sweden: 13.0mm rings for Mallard and Pochard, 7.0mm rings for Teal), so as to have similar resistance properties over time and hence similar recovery probabilities (Zimmerman et al., 2010; Souchay et al., 2014). In order for reward rings to be self- explicit (Pollock et al., 2001), those were labelled “RECOMPENSE/REWARD 70€ Tel. 00 33 4 90 97 27 87”. A 70€ reward was selected for this pilot study in order to reach a similar value to the $100 used in many former bird and fish reward mark reporting studies, considered to be over the value where mark reporting probability reaches close to 1 (e.g. Nichols et al., 1991, 1995; Scott et al., 2004; Royle & Garrettson, 2005; Meyer et al., 2012; Zimmerman et al., 2009).

A vast communication campaign was launched upon initiation of the programme, with the same article explaining the protocol and what to do with reward and regular rings being sent to all federal hunting magazines in France (all French hunters have to join a local hunting federation, of which there are ca. 90 in France, each generally having its own monthly magazine sent to hunters for free).

All recoveries were considered here, not only direct recoveries, defined as recoveries occurring the year of ringing. As opposed to some studies in North America, no check stations exist so that all recovered rings were voluntarily sent back by hunters, there were no solicited rings (see Arnold et al., 2020). Apart from one Teal and one Mallard, who were found freshly dead (likely from botulism), all recoveries were from hunting and reported by hunters quickly after recovery. Although all birds were initially ringed in France, recoveries could come from anywhere in the flyway. In practice, ca. 90% of the ringed birds (with or without a reward ring) were recovered in France (proportions and list of recovery countries in Table 1).

**Table 1.**
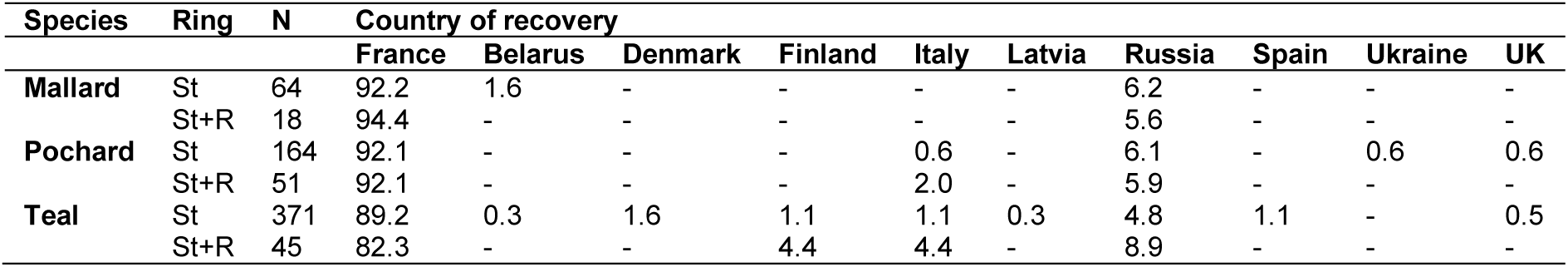
Total number of recoveries and relative percentages per country for each species with only a standard (St) or a standard plus a reward ring (St+R).

### Modelling

The definition of all terms is provided as a glossary in Supplementary Information Appendix A. We assumed that the reward value used in this study was high enough, so that the reporting probability of reward rings was 1. Thus, standard ring reporting probability (λ) can be directly estimated from recovery probabilities following Royle & Garrettson (2005): *λ = f Ú f’*, with *f* and *f’* the unconditional recovery probabilities for standard only and reward-ringed individuals, respectively. Hence, harvest probability (*h*) can be estimated following the relationship *h = f Ú λ* (Anderson & Burnham, 1976). Unconditional recovery probability (*f*) can be related to survival (*S*) and conditional recovery probability (*r*) with *f* = (1-*S*)\**r* (Gauthier & Lebreton, 2008), allowing to estimate the ring reporting rate as *λ = r Ú r’*.

Given that several parameters have to be estimated from other ones, we decided to use a Bayesian framework to propagate uncertainty over the estimation process. We used a Bayesian multi-state state-space model to estimate annual survival and conditional recovery probabilities following Schaub & Badia-Boher (2025), and integrating the direct estimation of the reporting probability and the derivation of the harvest probabilities for each species (Supplementary Information Appendix B and Figure S1).

The model proposed by Schaub & Badia-Boher allows for the joint use of all reencounters (live recaptures – including resighting, and dead recoveries). However, we decided to discard live resighting and recaptures to avoid complexity in the parameterization of reencounter probabilities, and we simply set recapture probabiliy to 0. Thus, our model is a Bayesian capture-mark-recovery model.

For each species, we included several effects on survival and recovery probabilities at the annual scale (Table 2): the age of the individual at ringing (AgeB: First-year of life *Juv* vs. After- First-Year individuals *Ad*), the ringing season (SeasB: *Fall* – August to October; *Winter* – November to January; *Spring* – February to March), time since marking (TSM: first year of marking *FY* vs. subsequent years *AFY*), presence of a reward ring (reward: standard only vs. standard + reward), and the presence of auxiliary marks for Teal only (saddle: with vs. without). We decided not to include time variation in our models to avoid issues of parameter identifiability, and ignored a sex effect given small sample sizes. Because not all individuals were ringed at the same moment throughout the year, parameters for the first year of ringing did not fully correspond to annual probabilities: survival probability for the first-year of ringing were indeed estimated over a 11- months interval until the next August 1st for individuals ringed in Fall, 7-months and 5-months intervals for individuals ringed in Winter and Spring, respectively. This is an inherent problem for waterfowl ringing in Europe, where breeding and moulting largely occurs in distant and inaccessible eastern regions, so a pre-hunting ringing season concentrated during few weeks in late summer, as done in North America, is not practical here.

**Table 2.** Parameterization of survival and recovery probabilities in the global model for each species.

| Parameter | Effect | Explanation | Species |
|---|---|---|---|
| **Survival** | *AgeB* | First-year and adult birds may have different survival probabilities | All |
|  | *SeasB* | Survival probability may differ between seasons of ringing | All |
|  | *TSM* | Ringing date and hence the length of the first event may affect survival probability estimate | All |
|  | *Saddle* | Nasal saddles may affect survival probability | Teal |
| **Recovery** | *TSM* | Direct and indirect recovery probabilities may differ | All |
|  | *Reward* | Reward rings should increase reporting by hunters | All |
|  | *Saddle* | Auxiliary marks may increase reporting by hunters | Teal |

Data management was performed and all Bayesian models were run using software R v.4.3.3 (R Core Team, 2024) and packages tidyverse and nimble (de Valpine et al., 2017; Wickham et al., 2019).

## Results

Detailed demographic parameter estimates for Mallard, Teal and Pochard are provided in Tables 3 and 4. Owing to asymmetrical distributions, mean and median values of estimates often differed, sometimes markedly, hence both are shown in the tables. Because of different ringing dates and hence differential exposure length to seasonal hunting, estimates often differed markedly between the first year of ringing and subsequent years: survival probability estimates were very high during the first year of ringing, including for birds ringed during fall (i.e. right during the hunting season) where it could reach as much as 0.78 ± 0.12 SD for unsaddled adult Teal and 0.77 ± 0.12 for juvenile Mallards up to the 1st of August of the next year (Table 3, Figure 2.A). Survival probabilities for birds ringed during winter or spring were even higher and corresponded to a period of the year with no hunting or close to the end of the hunting season (hence lower killing probabilities), and also correspond to shorter time periods until the next 1st of August. After that initial ringing year, annual survival probabilities (over 12 months proper) ranged from 0.39 ± 0.04 SD in saddled Teal (vs. 0.52 ± 0.04 SD in unsaddled Teal) to 0.68 ± 0.04 SD in Pochard (Figure 2B).

**Figure 2.**
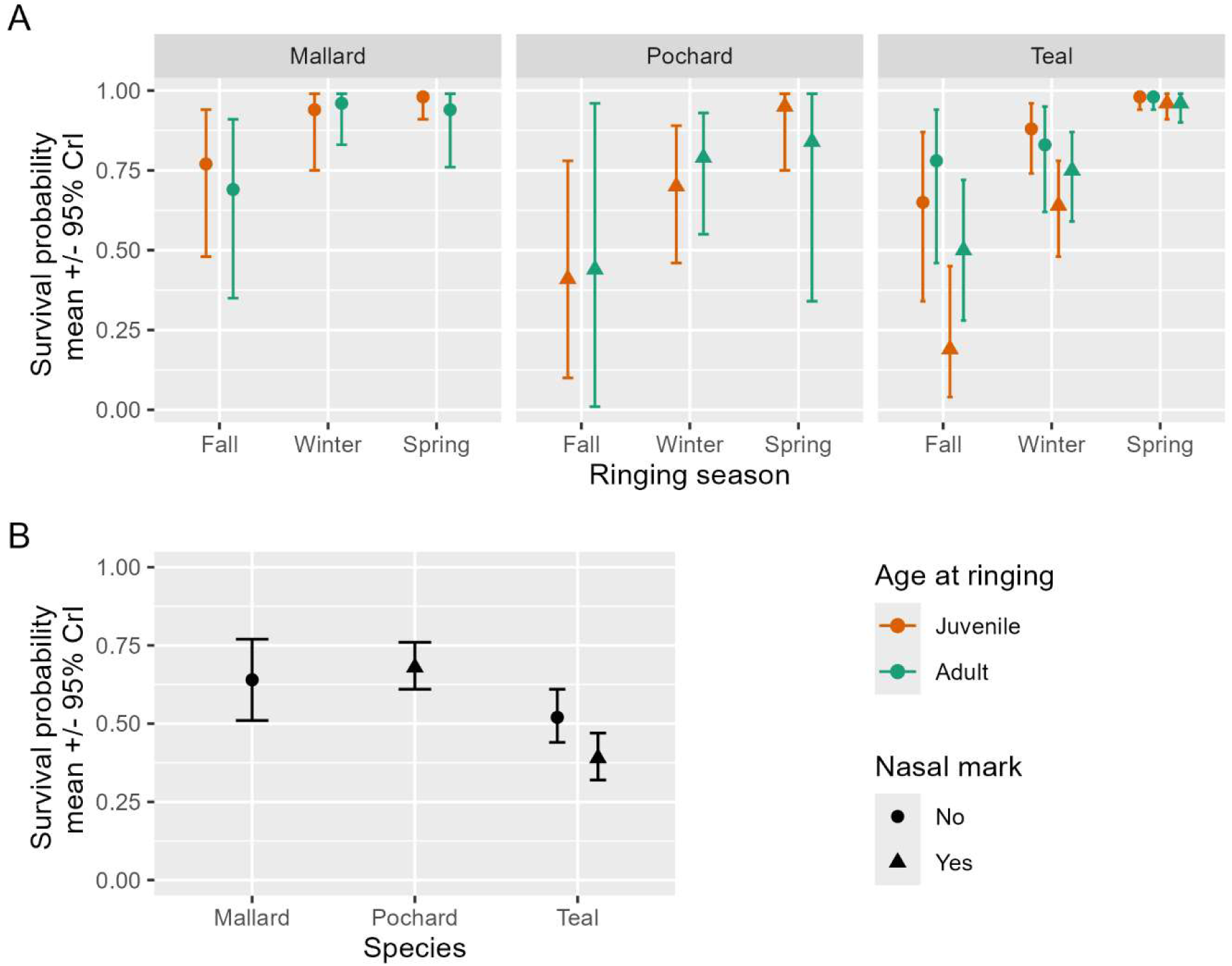
Survival probability estimate (estimated mean with 95% credible interval (CrI). **A**: Estimates for the first year of ringing, by species, ringing season, age at ringing (Juveniles: orange symbols; Adults: green symbols) and presence (triangle) or absence of a nasal saddle (dots). **B**: Estimates for subsequent years by species and presence of nasal saddle.

**Table 3.** Survival and conditional recovery probabilities estimated for each species and effect category. See Table 2 for a description of the effects. In addition to mean and standard deviation of the estimates, the quartiles and median value are also shown. Bold type indicates cases where mean and median values were markedly different.

| **Parameter** | **Species** | **Nasal saddle** | **Ringing age** | **Ringing season** | **Time Since Marking** | **Ringing type** | **Mean estimate** | **Standard deviation** | **2.5%** | **50%** | **97.5%** |
|---|---|---|---|---|---|---|---|---|---|---|---|
| **Survival**<br>***S*** | Mallard | No | Juv | Fall | FY | - | 0.77 | 0.12 | 0.48 | 0.79 | 0.94 |
|  |  | No | Ad | Fall | FY | - | 0.69 | 0.16 | 0.35 | 0.71 | 0.91 |
|  |  | No | Juv | Winter | FY | - | 0.94 | 0.07 | 0.75 | 0.97 | 0.99 |
|  |  | No | Ad | Winter | FY | - | 0.96 | 0.05 | 0.83 | 0.97 | 0.99 |
|  |  | No | Juv | Spring | FY | - | 0.98 | 0.02 | 0.91 | 0.98 | 0.99 |
|  |  | No | Ad | Spring | FY | - | 0.94 | 0.06 | 0.76 | 0.96 | 0.99 |
|  |  | No | - | - | AFY | - | 0.64 | 0.07 | 0.51 | 0.64 | 0.77 |
|  | Pochard | Yes | Juv | Fall | FY | - | 0.41 | 0.20 | 0.10 | 0.38 | 0.78 |
|  |  | Yes | Ad | Fall | FY | - | 0.44 | 0.34 | 0.01 | 0.43 | 0.96 |
|  |  | Yes | Juv | Winter | FY | - | 0.70 | 0.12 | 0.46 | 0.70 | 0.89 |
|  |  | Yes | Ad | Winter | FY | - | 0.79 | 0.10 | 0.55 | 0.80 | 0.93 |
|  |  | Yes | Juv | Spring | FY | - | 0.95 | 0.07 | 0.75 | 0.98 | 0.99 |
|  |  | Yes | Ad | Spring | FY | - | 0.84 | 0.17 | 0.34 | 0.91 | 0.99 |
|  |  | Yes | - | - | AFY | - | 0.68 | 0.04 | 0.61 | 0.68 | 0.76 |
|  | Teal | No | Juv | Fall | FY | - | 0.65 | 0.14 | 0.34 | 0.67 | 0.87 |
|  |  | Yes | Juv | Fall | FY | - | 0.19 | 0.11 | 0.04 | 0.17 | 0.45 |
|  |  | No | Ad | Fall | FY | - | 0.78 | 0.12 | 0.46 | 0.81 | 0.94 |
|  |  | Yes | Ad | Fall | FY | - | 0.50 | 0.11 | 0.28 | 0.51 | 0.72 |
|  |  | No | Juv | Winter | FY | - | 0.88 | 0.06 | 0.74 | 0.90 | 0.96 |
|  |  | Yes | Juv | Winter | FY | - | 0.64 | 0.08 | 0.48 | 0.64 | 0.78 |
|  |  | No | Ad | Winter | FY | - | 0.83 | 0.09 | 0.62 | 0.85 | 0.95 |
|  |  | Yes | Ad | Winter | FY | - | 0.75 | 0.07 | 0.59 | 0.75 | 0.87 |
|  |  | No | Juv | Spring | FY | - | 0.98 | 0.02 | 0.94 | 0.99 | 0.99 |
|  |  | Yes | Juv | Spring | FY | - | 0.96 | 0.02 | 0.91 | 0.97 | 0.99 |
|  |  | No | Ad | Spring | FY | - | 0.98 | 0.02 | 0.94 | 0.99 | 0.99 |
|  |  | Yes | Ad | Spring | FY | - | 0.96 | 0.02 | 0.90 | 0.96 | 0.99 |
|  |  | No | - | - | AFY | - | 0.52 | 0.04 | 0.44 | 0.52 | 0.61 |
|  |  | Yes | - | - | AFY | - | 0.39 | 0.04 | 0.32 | 0.39 | 0.47 |
| **Recovery**<br>***r*** | Mallard | No | - | - | Direct | St | 0.31 | 0.18 | 0.10 | 0.26 | 0.82 |
|  |  | No | - | - | Direct | St+R | 0.48 | 0.27 | 0.14 | 0.40 | 0.99 |
|  |  | No | - | - | Indirect | St | 0.08 | 0.02 | 0.05 | 0.08 | 0.12 |
|  |  | No | - | - | Indirect | St+R | 0.11 | 0.04 | 0.07 | 0.10 | 0.20 |
|  | Pochard | Yes | - | - | Direct | St | 0.39 | 0.20 | 0.17 | 0.32 | 0.91 |
|  |  | Yes | - | - | Direct | St+R | 0.49 | 0.24 | 0.21 | 0.41 | 0.99 |
|  |  | Yes | - | - | Indirect | St | 0.15 | 0.03 | 0.10 | 0.14 | 0.22 |
|  |  | Yes | - | - | Indirect | St+R | 0.22 | 0.06 | 0.13 | 0.21 | 0.36 |
|  | Teal | No | - | - | Direct | St | 0.23 | 0.11 | 0.09 | 0.20 | 0.49 |
|  |  | No | - | - | Direct | St+R | 0.61 | 0.27 | 0.16 | 0.59 | 0.99 |
|  |  | Yes | - | - | Direct | St | 0.23 | 0.05 | 0.16 | 0.22 | 0.34 |
|  |  | Yes | - | - | Direct | St+R | 0.26 | 0.07 | 0.17 | 0.24 | 0.42 |
|  |  | No | - | - | Indirect | St | 0.07 | 0.01 | 0.06 | 0.07 | 0.09 |
|  |  | No | - | - | Indirect | St+R | 0.11 | 0.03 | 0.07 | 0.11 | 0.19 |
|  |  | Yes | - | - | Indirect | St | 0.11 | 0.01 | 0.09 | 0.11 | 0.13 |
|  |  | Yes | - | - | Indirect | St+R | 0.15 | 0.04 | 0.10 | 0.14 | 0.24 |

**Table 4.** Reporting and harvest probabilities estimated for each species and effect category (considering a 100% reporting probability for reward-ringed individuals: only information from regular-ringed birds is shown). See Table 2 for a description of the effects. In addition to mean and standard deviation of the estimates, the quartiles and median value are also shown. Bold type indicates cases where mean and median values were markedly different.

| Parameter | Species | Nasal saddle | Ringing age | Ringing season | Time Since Marking | Ringing type | Mean estimate | Standard deviation | 2.5% | 50% | 97.5% |
|---|---|---|---|---|---|---|---|---|---|---|---|
| **Reporting**<br>λ | Mallard | No | - | - | FY | - | 0.70 | 0.22 | 0.31 | 0.70 | 0.99 |
|  |  | No | - | - | AFY | - | 0.76 | 0.19 | 0.37 | 0.79 | 0.99 |
|  | Pochard | Yes | - | - | FY | - | 0.81 | 0.14 | 0.52 | 0.83 | 0.99 |
|  |  | Yes | - | - | AFY | - | 0.70 | 0.16 | 0.43 | 0.68 | 0.99 |
|  | Teal | No | - | - | FY | - | 0.42 | 0.20 | 0.18 | 0.36 | 0.97 |
|  |  | Yes | - | - | FY | - | 0.90 | 0.11 | 0.60 | 0.94 | 0.99 |
|  |  | No | - | - | AFY | - | 0.70 | 0.19 | 0.37 | 0.68 | 0.99 |
|  |  | Yes | - | - | AFY | - | 0.76 | 0.17 | 0.43 | 0.77 | 0.99 |
| **Harvest**<br>***h*** | Mallard | No | Juv | Fall | FY | - | 0.09 | 0.04 | 0.03 | 0.08 | 0.19 |
|  |  | No | Ad | Fall | FY | - | 0.12 | 0.04 | 0.06 | 0.11 | 0.22 |
|  |  | No | Juv | Winter | FY | - | 0.02 | 0.02 | 0.00 | 0.01 | 0.07 |
|  |  | No | Ad | Winter | FY | - | 0.02 | 0.01 | 0.00 | 0.01 | 0.05 |
|  |  | No | Juv | Spring | FY | - | 0.01 | 0.01 | 0.00 | 0.01 | 0.03 |
|  |  | No | Ad | Spring | FY | - | 0.02 | 0.02 | 0.00 | 0.02 | 0.08 |
|  |  | No | - | - | AFY | - | 0.04 | 0.01 | 0.02 | 0.04 | 0.07 |
|  | Pochard | Yes | Juv | Fall | FY | - | 0.25 | 0.07 | 0.14 | 0.24 | 0.41 |
|  |  | Yes | Ad | Fall | FY | - | 0.21 | 0.11 | 0.03 | 0.21 | 0.44 |
|  |  | Yes | Juv | Winter | FY | - | 0.12 | 0.02 | 0.09 | 0.12 | 0.18 |
|  |  | Yes | Ad | Winter | FY | - | 0.09 | 0.02 | 0.05 | 0.08 | 0.14 |
|  |  | Yes | Juv | Spring | FY | - | 0.02 | 0.02 | 0.00 | 0.01 | 0.09 |
|  |  | Yes | Ad | Spring | FY | - | 0.06 | 0.05 | 0.00 | 0.04 | 0.20 |
|  |  | Yes | - | - | AFY | - | 0.07 | 0.02 | 0.04 | 0.07 | 0.12 |
|  | Teal | No | Juv | Fall | FY | - | 0.19 | 0.08 | 0.06 | 0.18 | 0.37 |
|  |  | Yes | Juv | Fall | FY | - | 0.21 | 0.04 | 0.14 | 0.20 | 0.31 |
|  |  | No | Ad | Fall | FY | - | 0.11 | 0.05 | 0.04 | 0.11 | 0.24 |
|  |  | Yes | Ad | Fall | FY | - | 0.12 | 0.03 | 0.08 | 0.12 | 0.19 |
|  |  | No | Juv | Winter | FY | - | 0.06 | 0.03 | 0.02 | 0.06 | 0.12 |
|  |  | Yes | Juv | Winter | FY | - | 0.09 | 0.02 | 0.06 | 0.09 | 0.13 |
|  |  | No | Ad | Winter | FY | - | 0.09 | 0.04 | 0.03 | 0.08 | 0.18 |
|  |  | Yes | Ad | Winter | FY | - | 0.06 | 0.02 | 0.04 | 0.06 | 0.10 |
|  |  | No | Juv | Spring | FY | - | 0.01 | 0.01 | 0.00 | 0.01 | 0.03 |
|  |  | Yes | Juv | Spring | FY | - | 0.01 | 0.01 | 0.00 | 0.01 | 0.02 |
|  |  | No | Ad | Spring | FY | - | 0.01 | 0.01 | 0.00 | 0.01 | 0.03 |
|  |  | Yes | Ad | Spring | FY | - | 0.01 | 0.01 | 0.00 | 0.01 | 0.02 |
|  |  | No | - | - | AFY | - | 0.05 | 0.02 | 0.03 | 0.05 | 0.09 |
|  |  | Yes | - | - | AFY | - | 0.09 | 0.02 | 0.06 | 0.08 | 0.15 |

Conditional recovery probability estimates varied among species, with higher values for direct than for indirect recoveries (Fig. 3). The presence of a reward ring increased the recovery probability for each case except for saddled Teal, which had similar direct recovery probabilities with or without a reward ring (0.23 vs. 0.26 for standard and reward rings, respectively). Regarding values for standard rings only, direct recovery probabilities ranged from 0.23 for Teal to 0.39 ± 0.20 for Pochard, while indirect recovery probabilities ranged from 0.07 ± 0.01 for unsaddled Teal to 0.15 ± 0.03 for Pochard (Table 3).

**Figure 3.**
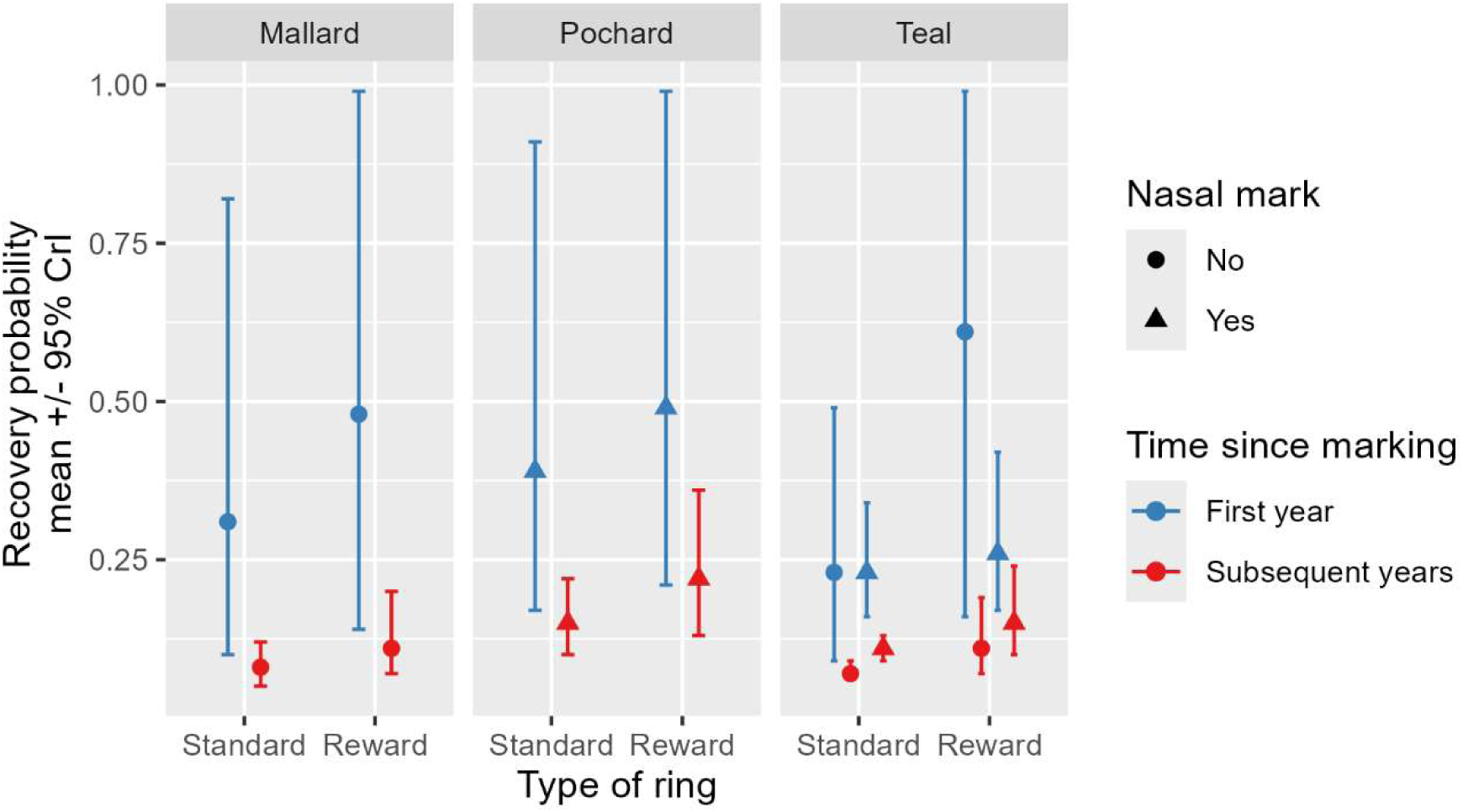
Recovery probability estimates by species, type of ring (standard vs. reward + standard rings), time since marking (blue vs. red), and presence (triangle) or absence of a nasal saddle (dots).

Reporting probability for regular rings (assuming a 100% reporting probability of reward rings) was particularly high during the year of ringing for birds wearing a nasal saddle (0.81 ± 0.14 SD in Pochard, 0.90 ± 0.11 in saddled Teal) and, conversely, very low in unsaddled Teal during that year (0.42 ± 0.20 SD, Fig. 4). In all other cases (Mallard during the year of ringing and all three species during later years) the reporting probability was very similar, ranging from 0.70 ± 0.19 SD in unsaddled Teal (0.76 ± 0.17 SD in saddled Teal) to 0.76 ± 0.19 SD in Mallard (Table 4).

**Figure 4.**
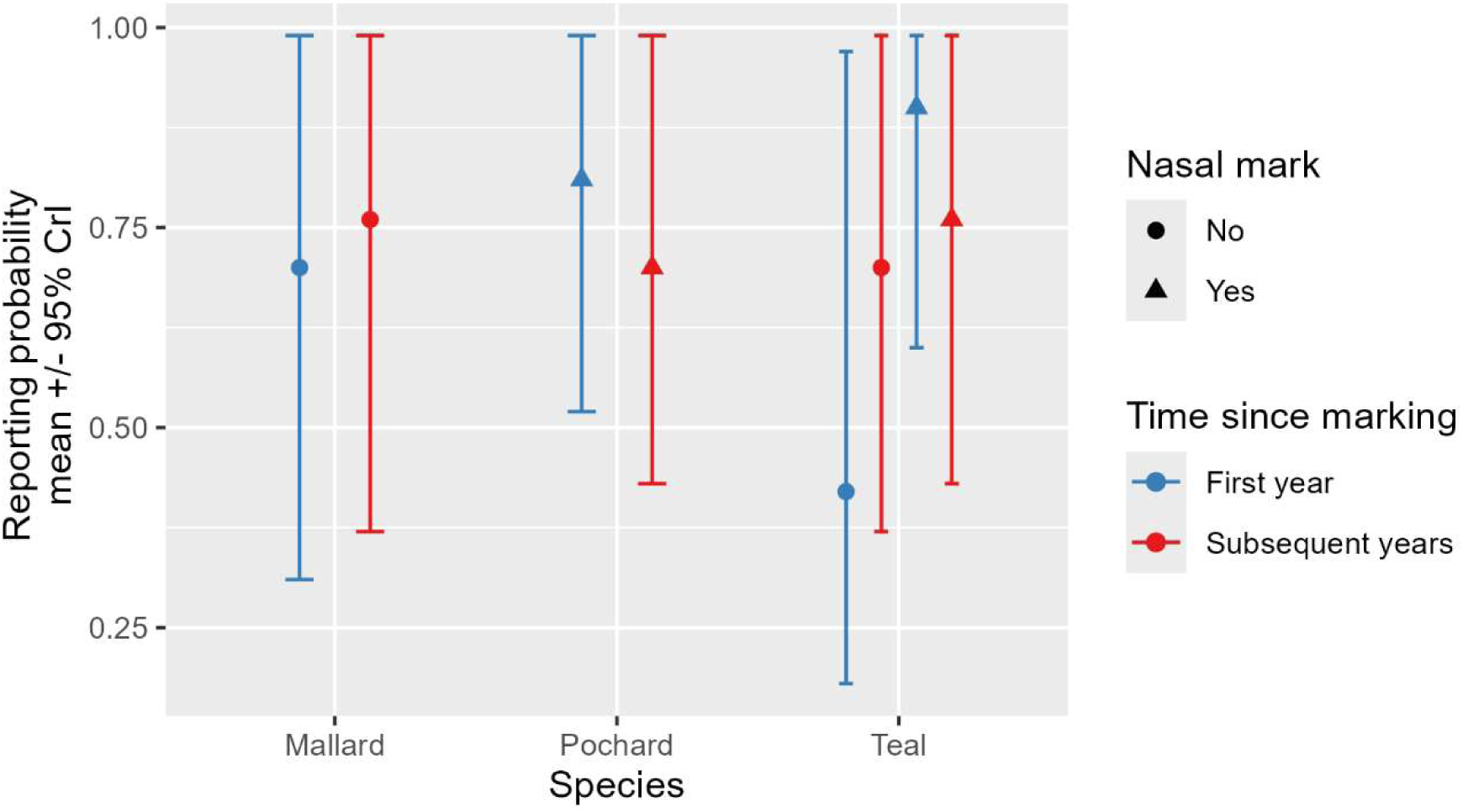
Reporting probability estimates by species, time since marking (first year [blue] vs. subsequent years [red]), and presence (triangle)/ absence (dot) of a nasal mark.

From the results above it was possible to estimate harvest probabilities for the various species and effect categories (Table 4 and Fig. 5). Although it covered only a part of a year (since birds could be ringed from August to March), annual harvest probability was always greater for the year of ringing (when it could be as high as 0.25 ± 0.07 SD in juvenile Pochard ringed during fall) than for later years, when it never exceeded 0.09 ± 0.02 SD in Teal wearing a nasal saddle. During the year of ringing, harvest probability declined from birds ringed in fall, to those ringed in winter then spring, according to the fact that they were exposed to fewer months of hunting, respectively. Juveniles ringed during fall tended to experience a greater harvest probability than adults during the first year of ringing, but such a difference faded over the seasons of that first year as harvest probability plummeted from fall-ringed to spring-ringed. Harvest probability of saddled and unsaddled Teal was always similar during the first year of ringing, whatever their ringing age or season. During later years, nasal saddled Teal were harvested to a greater extent than their unsaddled conspecifics (0.09 ± 0.02 SD and 0.05 ± 0.02 SD, respectively, Table 4).

**Figure 5.**
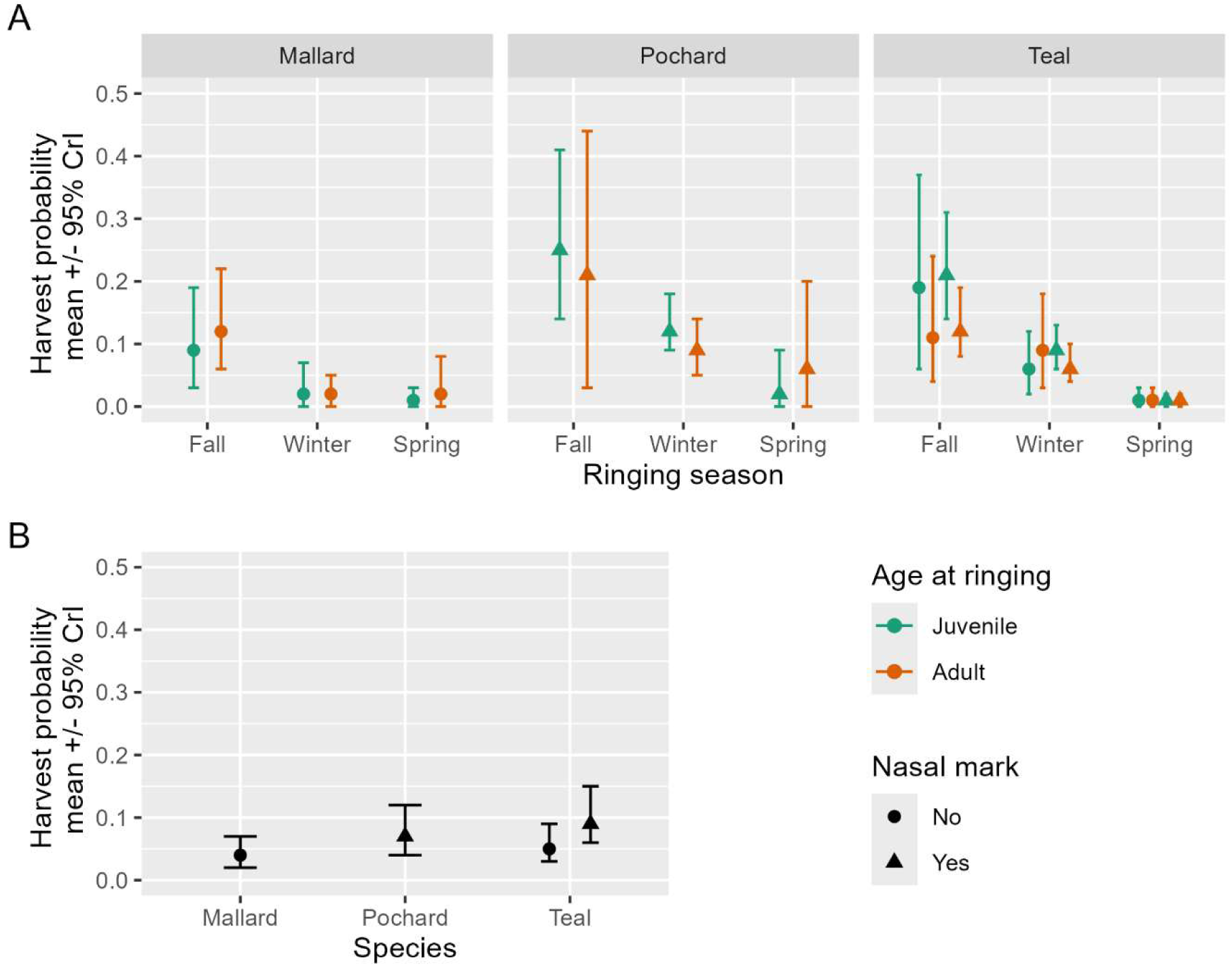
Harvest probability estimate (estimated mean with 95% credible interval (CrI). **A**: Estimates for the first year of ringing, by species, ringing season, age at ringing (Juveniles: orange symbols; Adults: green symbols) and presence (triangle)or absence of a nasal saddle (dots). **B**: Estimates for subsequent years by species and presence of nasal saddle.

## Discussion

The results presented here provide the first ever estimates of ring reporting probabilities and therefore harvest probabilities from capture-mark-recovery for waterbirds in Europe. People (notably hunters, since most ring recoveries came from shot birds) apparently report found rings at a high 3/4 ratio. Ducks captured early in the year and exposed to most of the hunting season suffer high harvest probabilities during their first year of ringing, a probability falling to 4-9% annually during later years. Given that all the birds used here are ringed in France, this result is consistent with an earlier study showing a greater hunting mortality within France compared to other countries for birds ringed in Camargue, southern France (Plaquin et al., 2025).

It should be kept in mind that the Bayesian approach used here means that all parameters are simultaneously estimated and informing each other, and that sample sizes were sometimes limited for some effect categories (Supplementary information Table S1), notably during the year of ringing, causing uncertainty that is propagated from a parameter to the others.

Nevertheless, the results obtained for annual survival probabilities are very consistent with the literature, especially when considering survival during years after the initial year of ringing, which is less affected by the sample size issues mentioned above and do cover a full 12-months period: the 0.52 value for unsaddled Teal is similar to the results obtained for Green-winged Teal (*Anas carolinensis*) in North America by Chu et al. (1995; 0.51 in females, 0.55 in males), Devineau et al. (2010; 0.54 both sexes combined), Fleming & Howell (2013; 0.50 in adult females and 0.58 in adult males) or Thompson et al. (2022; 0.44 to 0.49 in adult females, 0.56 to 0.58 in adult males), but lower than results of Lake et al. (2006) for adult birds (0.64 in males; 0.66 in females). The present results are also consistent with the few earlier data available from Europe for Eurasian Teal, i.e. around 0.55 in adults depending on UK ringing sites in Gitay et al. (1990) or 0.49 both sexes combined in Devineau et al. (2010) for Camargue-wintering birds.

Survival probability estimates for Pochard are scarce, too, but the 0.68 annual probability obtained here for years after the year of ringing aligns with the 0.71 value of Kharitonov (2017), and is only slightly higher than the 0.65 for adult females in Blums et al. (1996), from the specific case of Engure Lake in Latvia, where a heavy predator-control program was in place. It is however significantly lower than the estimates for the UK and Switzerland in Folliot et al. (2020), where female values were 0.67 and 0.69 but male values were 0.81 and 0.75, respectively. It is possible that hunting pressure over these birds was lower in those countries, notably in the UK where recovery probabilities were notably much lower than those reported in France (respectively 0.04-0.05 against 0.18 for direct recoveries and ca. 0.09 against 0.15 for indirect recoveries, see Folliot et al. 2020). Moreover, a large proportion of the Pochards ringed on our Grand-lieu study site during Fall and Winter would be resident (Gourlay-Larour et al., 2014) and hence would always suffer a higher hunting pressure. Indeed, 90 to 95% of hunting recoveries from such birds were done in France.

Given their extremely wide distribution and the fact that some populations are composed of both wild, farmed, and hybrid individuals (Champagnon et al., 2023), Mallards around the world experience a wide variety of environments and hence show a large range of demographic values. During years after the year of ringing, annual survival probability of Mallard here was 0.64, which nonetheless matches European ringing results of Söderquist et al. (2021) from Sweden (0.64 in females, 0.71 in males) or Tavecchia et al. (2001) from southern France (0.61 in females, 0.66 in males), North American estimates such as in Giudice (2003; 0.56 in juvenile males to 0.66 in adult males), Devers et al. (2021; 0.49 in juvenile females to 0.63 in adult males), or the New Zealand values from McDougall & Amundson (2017; 0.49 in juvenile females to 0.66 in adult males).

Despite the caveats mentioned above for sample size in some categories, differences in survival probabilities between effect categories recorded here are also consistent with available knowledge (review in Nicolai et al. 2026). Survival probability during the year of ringing was generally higher for birds ringed as adults than juveniles (Kear, 2005; Baldassarre, 2014), higher for the larger species (Pochard) and lower for the smaller one (Teal), in line with earlier waterfowl knowledge and life-history theory (Boyd, 1962; Stearns, 1992; Koons et al., 2014). The addition of a visual mark (nasal saddle) led to a lower survival probability in Teal (Figure 3), especially in juveniles, but a higher reporting probability. Such reduced survival of nasal marked ducks echoes earlier work (Deane et al., 2021; Guillemain et al., 2026; Guillemain et al., in prep.). The presence of the saddle may also question hunters and lead them to report saddled birds more, with the interplay between both probabilities making interpretation more difficult (see also Reinecke et al., 1992; Caswell et al., 2012; Arnold et al., 2016; Szymanski et al., 2020). Survival probability during the year of ringing increased from birds ringed in fall (at the beginning of the hunting season) to those ringed in spring (after the hunting season). This is consistent with the fact that ducks ringed later have to survive fewer months until the beginning of the next occasion as we coded it (i.e., next 1st of August), hence partly a modelling artefact. The present results however also support the idea that hunting mortality is at least partially additive to natural mortality in these species (Cooch et al., 2014), since a longer exposure to the hunting season during the year of ringing (e.g. autumn-ringed versus late winter-ringed birds) caused a much lower survival rate associated with a much greater harvest rate. Inter-individual heterogeneity in vulnerabiity to harvest may nevertheless lead to some sort of partial compensation (Grzegorczyk et al., in prep).

A quarter to two-thirds of the fitted rings were recovered during the year of ringing, which is consistent with the low annual survival probabilities in these species, and the fact that mortality is most often human-related in such huntable species (hence leading to greater recovery probabilities compared to deaths resulting from natural causes). The presence of a reward ring often doubled such conditional or ‘Seber’ recovery probability (Gauthier & Lebreton, 2008): reward rings hence led to a greater probability to get rings back, independently of mortality probability, indicating that the reward experiment was effective in boosting the likelihood of people sending rings back when there was a reward. The fact that the presence of a saddle may be associated with a slightly higher ring recovery probability among reward-ringed Teal after the year of ringing (i.e., indirect recoveries) may however suggest that the 70€ reward may not systematically lead to a 100% reporting probability, but the presence of a saddle may have caused curiosity leading to a further little increase in reporting (e.g., Henny & Burnham, 1976; Evrard, 1996). It should also be noted that we only used two modalities here, which can be considered as a pilot study: regular rings with no rewards and reward rings all with a 70€ amount. Other papers have sometimes used different reward values, relying on a regression analysis to estimate reporting probability of regular rings (e.g. Royle & Garrettson, 2005). We advocate the continuation of reward ring schemes to evaluate possible changes in reporting and harvest probabilities in the future, and to expand such schemes to other hunted species. Future studies should also aim to evaluate the current reward value required in Europe to obtain a 100% reporting probability, hence the most efficient use of public money. Future studies should also aim for sample sizes large enough to allow the consideration of a sex effect in the analyses, since it is very likely that the breeding risks only incurred by females leads to a lower survival probability in adults of this sex (see references above).

The present analysis however provides the first ever estimate of ring reporting probability for waterfowl in Europe, and such estimates were very similar among species beyond the year of ringing (indirect recoveries and reports), ranging from 0.70 to 0.76. Such a lack of differences among species is consistent with the results obtained in North America (for species different than those studied here) by Garretson et al. (2014) and Arnold et al. (2020). The fact that three fourths of the regular rings found by people are reported is very good news, and shows confidence of the public in the ringing process, the scientists and managers using these data (see also Guillemain, 2010). This is not so surprising given the long-standing relationships and outreach efforts of Office Français de la Biodiversité towards the hunting community in particular, with regular information in hunting magazines and conferences about what the ringing data are used for, helping to build trust. Results for the year of ringing, and notably the very low reporting probability in unsaddled Teal (0.42) and the very high probability in saddled Teal (0.90), do illustrate the curiosity effect caused by the saddles described above, but could have been amplified by small sample sizes. The reporting probabilities estimated here for years after the year of ringing are in the range of current probabilities for North American Mallard (0.67 to 0.81 depending on flyways, Boomer et al., 2013) or common North American waterfowl species (ca. 0.80 after Arnold et al., 2020), but the later increased sharply after rings were engraved with a toll-free number or web address on the rings at the end of the 1990s (Royle & Garrettson, 2005; Garrettson et al., 2014; Arnold et al. 2020). This suggests the present results for Europe, although already quite satisfactory, could easily be improved by more information on the rings.

Knowledge of the ring reporting probabilities allows computation of harvest probabilities in the three species, which had never been done so far in European waterfowl. The values obtained here for the first year after ringing are high in some categories of individuals ringed during fall, especially in Teal where it ranged 0.11-0.21 and in Pochard where it ranged 0.21-0.25, while this is for less than 12 months until the beginning of the next occasion on the following 1st of August (hence a proper 12-month annual estimate would be even higher). As a comparison, annual harvest probability of Green-winged Teal in North America was 0.07 after Devineau et al. (2010; considering a ring reporting probability of 30%, hence this was likely overestimated), ranged 0.07 to 0.13 after Flemming & Howell (2013) and 0.05 to 0.14 depending on age and sex classes in Thompson et al. (2022). In North America, based on known demographic processes of the different species, Johnson et al. (2019) evaluated that harvest probability should range, depending on whether hunting policy aims at being conservative to liberal, from 0.06 to 0.12 in Green-winged Teal, 0.08 to 0.16 in Mallard and 0.06 to 0.13 in ring-necked duck (*Aythya collaris*), a diving duck like Pochard. The expected harvest probability under liberal hunting seasons is currently estimated at 0.11-0.14 for adult male Mallard and 0.12 for Green-winged Teal (U.S. Fish and Wildlife Service, 2021). These values are generally higher than the estimates obtained here beyond the year of ringing, which correspond better to North American harvest probabilities under a restrictive regulation (0.05 to 0.07 in Mallard, U.S. Fish and Wildlife Service, 2021). These results suggest ducks ringed in France do encounter a relatively high hunting pressure throughout Europe during the first months after ringing, but are subsequently relatively little exposed during later years (maybe because of initial selection and learning to avoid hunted areas). There are no earlier European values which would allow assessing how duck harvest probability has changed in Europe. From Teal ringed between 1954 and 1976 in Southern France, Devineau et al. (2010) estimated a mean 0.227 harvest probability considering a 0.32 reporting probability. If the reporting probability was then already rather around 0.70 as in modern years here, historic harvest probability would actually have been 0.10, which is relatively similar to the current value.

Two of the species considered here, Pochard and Teal, are scrutinized by the European Commission owing to declining breeding populations in the European Union (European Environment Agency, 2020; link to the data on page 26). A former analysis suggested that EU hunting harvest was not sustainable in these populations (Carboneras et al., 2024), leading the Commission to call for more restrictive regulation. As a consequence, hunting quotas or moratoriums have recently been introduced in several countries, especially for Pochard, with a call for the implementation of population dynamics models. Such models however relied on strong assumptions about ring reporting rates and hence harvest probabilities, often using dated North American value, generally from a different species (i.e. Mallard). The present results will now be useful to parameterize such models with proper modern European estimates. Knowledge of harvest probability will also help disentangle natural and hunting mortality sources, hence inform the position of species along the additive/compensatory gradient, which is a prerequisite to establish and monitor the efficiency of harvest regulations. For Common Pochard and Teal specifically, the present value can be taken as a benchmark corresponding to a period without any quota, allowing to evaluate the effects of future harvest policy within an adaptive management framework.

## Acknowledgements

We would like to thank Kathy Fleming (US Fish and Wildlife Service) for discussions about ring reporting probability in (Green-winged) Teal, Jim Nichols (USGS) for technical advice while setting up the programme, and Jean-Marie Boutin for convincing technical, administrative and financial authorities that deploying reward rings in France would be a good idea. Comments by Jeffrey Hostetler, Céline Bellard and Todd Arnold greatly helped revising the initial manuscript. A preprint version of this article has been peer-reviewed and recommended by PCI Ecology (https://doi.org/10.24072/pci.ecology.101031).

## Conflict of interest disclosure

The authors declare that they comply with the PCI rule of having no financial conflicts of interest in relation to the content of the article.

### Data, scripts, code, and supplementary information availability

Data, scripts and supplementary information are available online (https://doi.org/10.5281/zenodo.23318764; Souchay et al, 2026).

